# Early sensory deprivation drives local reorganization of sensory integration within a conserved global hierarchy

**DOI:** 10.64898/2026.06.14.732169

**Authors:** Wei Wei, Annahita Sarré, Sami Abboud, Francesco Alberti, R. Austin Benn, Robert Scholz, Victoria Shevchenko, Alexander Holmes, Ulysse Klatzmann, Tamara Vanderwal, Elizabeth Jefferies, Marcin Szwed, Olivier Collignon, Laurent Cohen, Daniel S. Margulies

## Abstract

The human brain processes sensory information through a hierarchical system, from primary to higher-level regions, integrating inputs across modalities to support perception and cognition. While early sensory loss triggers widespread neuroplastic changes, its impact on integration across the cortical hierarchy remains unclear. Here, we examined the cortical reorganization of individuals with early blindness and deafness using a sensory integration framework that quantifies how brain regions prioritize different sensory inputs across the hierarchy. We found that early sensory deprivation drives highly localized reorganization adjacent to the deprived primary cortical areas: extrastriate cortex in early blindness and the superior temporal cortex in early deafness. These findings were further corroborated by analysis of the functional gradients, which found reorganization within these sensory regions. Notably, the hierarchy was largely preserved across groups. However, the sensory integration framework uniquely detected reorganization in language-related regions in deaf individuals with knowledge of a visual communication system known as cued speech. The specific differences between early deaf and hearing individuals remained restricted to superior temporal cortex. Together, our findings demonstrate that early sensory deprivation drives targeted reorganization adjacent to the affected primary sensory cortex, while preserving the overall hierarchy of cortical integration.

## Introduction

The human brain integrates information across sensory modalities to support perception and cognition (Jung et al., 2022; Spence & Driver, 2004). This integration relies on a hierarchical cortical organization, in which sensory information progresses from primary to higher-order regions, enabling coordinated interactions across sensory modalities. When one sensory modality is lost early in life, these interactions are disrupted, triggering widespread neuroplastic changes across the cortex (Bavelier & Neville, 2002; Merabet & Pascual-Leone, 2010; Rauschecker, 1995; Ricciardi & Pietrini, 2011).

The absence of early sensory experience can lead to large-scale functional reorganization. A hallmark of this reorganization is cross-modal plasticity, which refers to the reassignment of deprived cortical territories to process inputs from remaining modalities. For example, the occipital cortex of blind individuals responds to auditory and tactile stimulation (Arno et al. 2001; Amedi et al. 2007; Gougoux et al. 2005; Poirier et al. 2006; Collignon et al. 2007; Voss et al. 2008; Merabet et al. 2008; Uhl et al. 1991; Sadato et al. 1996), while the superior temporal cortex (STC) of early deaf individuals responds to visual and somatosensory information (Finney et al. 2001; Finney et al. 2003; MacSweeney et al. 2002; Fine et al. 2005; Auer et al. 2007; Emmorey et al. 2011; Leonard et al. 2012; Bottari et al. 2014; Karns et al. 2012; Cardin et al. 2013; Bola et al. 2017; Zimmermann et al. 2021). To explain the crossmodal adaptations, early accounts emphasized sensory substitution, proposing that the deprived cortex is reassigned to process alternative sensory inputs. This perspective has been refined by the supramodal/metamodal theory, which posits that cortical regions are organized according to the computations they perform rather than the modality of their inputs (Ricciardi et al., 2007; Amedi et al., 2010; Dormal & Collignon, 2011). Consistent with this view, deprived sensory cortex often retains functional selectivity resembling that observed under typical sensory experience. In early blindness, occipital regions exhibit selectivity for words (Reich et al., 2011; Matuszewski et al., 2025), motion (Poirier et al., 2004; Dormal et al., 2016), places (Wolbers et al., 2011; He et al., 2013), bodies (Kitada et al., 2014; Striem-Amit & Amedi, 2014), tools (Peelen et al., 2013), and shapes (Amedi et al., 2007; Xu et al., 2023). Similarly, in congenitally deaf individuals, auditory regions selective for voices or motion become tuned to faces or visual motion (Benetti et al., 2017, 2021), and posterior temporal regions preserve sensitivity to frequency-related perceptual dimensions across modalities (Bola et al., 2017). These findings suggest that early deprivation may preserve canonical computational roles, raising the possibility that cross-modal plasticity operates within a largely intact functional architecture. However, it remains unclear how such modality-specific substitution reshapes the intrinsic organization within the deprived cortex and beyond.

Notably, deprived sensory areas also contribute to higher-order cognitive domains. In early blindness, the occipital cortex participates in working memory (Amedi et al., 2003; Bonino et al., 2008; Park et al., 2011; Rimmele et al., 2019), numerical processing (Kanjlia et al. 2016; Crollen et al. 2019), and language (Röder et al. 2002; Amedi et al. 2004; Bedny et al. 2011; Watkins et al. 2012; Abboud and Cohen 2019). Likewise, auditory regions in early deaf individuals are recruited during visual working memory (Cardin et al., 2018; Ding et al., 2015). In this context, these observations challenge a strictly computation-preserving account, as the recruitment of deprived sensory regions for higher-order cognitive functions extends beyond their canonical sensory computations, and raise a central unresolved question: Does early sensory deprivation alter how sensory information is integrated along the cortical hierarchy, or does reorganization occur within a preserved macroscale functional architecture?

To address these questions, we employed a multisensory analytic framework that characterizes both the intrinsic nature of sensory processing and the macroscale functional architecture of the cortex. First, we utilized our recently developed function-based sensory integration model to quantify the reorganization of sensory integration strategies across the cortical hierarchy (Wei et al., 2024). This model decomposes neural activity into two distinct dimensions: sensory magnitude, which indexes the level of sensory dependence versus abstract integration; and sensory angle, which captures the proportional contribution of visual, auditory, and somatosensory modalities. Together, these metrics allow us to pinpoint how the integration landscape shifts within the sensory-modality deprived brain. Second, to investigate whether this reorganization disrupts the brain’s functional connectivity pattern, we used gradient analysis to characterize axes of functional organization. Gradients represent the most prominent functional axes of the connectome in low-dimensional manifolds, thus providing a data-driven coordinate representation of brain connectivity. By integrating these two approaches and applying them to the resting-state fMRI data acquired from early blind and early deaf individuals, we aimed to determine whether the changes in sensory integration observed in deprived individuals occur within a preserved functional architecture, or whether they result in a remapping of the global cortical hierarchy.

## Methods

### Data overview

This study comprises four previously published datasets: three concerning early blindness and one concerning early deafness (Table 1). Each early blind dataset includes one group of early blind individuals and one group of sighted controls. The three early blind datasets were collected in Montreal (Canada), Trento (Italy), and Paris (France). The majority of blind participants were blind from birth, with several blind within months of birth.

**Table 1.** Demographic information.

| Table 1. Demographic information |  |  |  |  |  |  |
| --- | --- | --- | --- | --- | --- | --- |
| Study | Site | Group | Age<br>(mean ± SD) | Sex | between-group<br>difference (EB vs SC) | between-site<br>difference |
| early<br>blind | Montreal | EB | 51 ± 14.3 | 7F, 10M | age: $t = -0.24$, $p = 0.81$<br>sex: $\chi^2 = 0$, $p = 1$ | age:<br>$f = 29.99$,<br>$p = 8.03e-11$<br>sex:<br>$\chi^2 = 1.86$,<br>$p = 0.39$ |
|  |  | SC | 52.1 ± 12.4 | 8F, 10M |  |  |
| | Trento | EB | 32.8 ± 4.6 | 10F, 6M | age: $t = 0.3$, $p = 0.77$<br>sex: $\chi^2 = 0.1$, $p = 0.76$ | |
|  |  | SC | 32.2 ± 7.7 | 12F, 11M |  |  |
| | Paris | EB | 45.5 ± 10.9 | 6F, 5M | age: $t = 0.55$, $p = 0.59$<br>sex: $\chi^2 = 0$, $p = 1$ | |
|  |  | SC | 42.8 ± 13.1 | 8F, 5M |  |  |
| early<br>deaf | Paris | Deaf | 37.7 ± 7.9 | 12F, 7M | age: $f = 0.3$, $p = 0.74$<br>sex: $\chi^2 = 0.94$, $p = 0.63$ | |
|  |  | Hear | 38.8 ± 11.5 | 16F, 5M |  |  |
|  |  | Ctrl | 36.3 ± 10.7 | 13F, 7M |  |  |

*EB, early blind; SC, sighted control; F, female; M, male; Deaf, early deaf with cued speech; Hear, hearing with cued speech; Ctrl, hearing with no cued speech.*

The early deaf dataset, acquired in Paris, includes three groups, in order to disentangle the influences of deafness and of language experience, which are both factors in the functional reorganization observed in deaf individuals. Deaf participants communicated in the French language, with the support of Cued Speech (CS), a system that complements lip reading and conveys the full phonological content of speech through a combination of lip movements and manual cues. To dissociate the effects of auditory deprivation from those of CS-based language exposure, the early deaf dataset includes: a group of early deaf individuals with knowledge of CS (Deaf), a group of hearing controls with knowledge of CS (Hear), and a group of hearing controls with no knowledge of CS (Ctrl).

Demographic information for all four datasets is presented in Table 1.

### MRI data

The Montreal dataset was acquired on a 3T Siemens MAGNETOM Prisma fit scanner equipped with a 32-channel head–neck coil. Resting-state functional images were collected using a simultaneous multislice echo-planar imaging sequence, with slices oriented parallel to the bicommissural plane and an anterior-to-posterior phase-encoding direction (TR = 785 ms, TE = 30 ms, flip angle = 54°, multiband factor = 3). Images were acquired at 3 mm isotropic resolution (field of view = 192 × 192 mm, matrix = 64 × 64, 42 axial slices). High-resolution T1-weighted anatomical images were acquired using a magnetization-prepared rapid gradient-echo sequence (TR = 2300 ms, TE = 2.26 ms, inversion time = 900 ms, flip angle = 8°, field of view = 288 × 288 mm, matrix = 256 × 256, 176 contiguous sagittal slices), yielding 1 mm isotropic voxels. Recruitment and experimental procedures were approved by the Multicentric Research Ethics Board of the Regroupement Neuroimagerie du Québec.

The Trento early blind dataset was acquired at the Center for Mind/Brain Sciences, University of Trento (Italy), using a 3T Siemens MAGNETOM Prisma scanner equipped with a 64-channel head–neck coil. Functional images were collected using a simultaneous multislice echo-planar imaging sequence, with slices oriented parallel to the bicommissural plane and an anterior-to-posterior phase-encoding direction (TR = 1000 ms, TE = 28 ms, flip angle = 59°, multiband factor = 5). For the early blind group and the main sighted control group, images were acquired at 3 mm resolution (field of view = 198 × 198 mm, matrix = 66 × 66, 65 axial slices, slice thickness = 3 mm with a 0.3 mm inter-slice gap). For a subset of nine participants from an independent sighted group, higher-resolution functional images were acquired at 2 mm resolution (field of view = 200 × 200 mm, matrix = 100 × 100, 65 axial slices, slice thickness = 2 mm with a 0.2 mm gap). T1-weighted anatomical images were acquired using a magnetization-prepared rapid gradient-echo sequence (TR = 2140 ms, TE = 2.9 ms, inversion time = 950 ms, flip angle = 12°, field of view = 288 × 288 mm, matrix = 288 × 288, 208 contiguous sagittal slices), yielding 1 mm isotropic voxels. The study was approved by the Ethics Committee of the University of Trento.

The Paris early blind dataset was acquired on a 3T Siemens MAGNETOM Verio scanner equipped with a 32-channel head coil. Resting-state functional images were collected using a single-echo echo-planar imaging sequence (TR = 2500 ms, TE = 30 ms, flip angle = 80°), with a slice thickness of 3 mm and whole-brain coverage. High-resolution T1-weighted anatomical images were acquired using a magnetization-prepared rapid gradient-echo sequence (TR = 2300 ms, TE = 3.1 ms, flip angle = 9°), with a slice thickness of 0.8 mm. Recruitment and experimental procedures were approved by the local ethics committee (Abboud & Cohen, 2019).

The Paris early deaf dataset was acquired on a 3T Siemens MAGNETOM Prisma scanner equipped with a 64-channel phased-array head coil. Resting-state functional images were collected using a multi-echo echo-planar imaging sequence (TR = 1660 ms, TEs = 14.2/35.39/56.58 ms, flip angle = 74°), with a slice thickness of 2.5 mm and whole-brain coverage. High-resolution T1-weighted anatomical images were acquired using a magnetization-prepared rapid gradient-echo sequence (TR = 2300 ms, TE = 2.76 ms, flip angle = 9°), yielding 1 mm isotropic voxels. The study was approved by the Comité de Protection des Personnes Est-III (CPP No. 20.11.05) (Sarré & Cohen, 2025).

### Preprocessing

The early blind and early deaf datasets were analyzed together due to their shared focus on cortical reorganization following early sensory deprivation. However, the datasets differed in fMRI acquisition protocols, most notably the use of single-echo versus multi-echo imaging, and therefore required dataset-specific preprocessing pipelines.

The early blind datasets were preprocessed using Micapipe (Cruces et al., 2022), a surface-based pipeline optimized for accurate cortical alignment and parcellation. This approach was selected to ensure precise localization of primary sensory cortices and reliable surface correspondence across individuals, which is critical for mapping cross-modal functional connectivity in populations with altered sensory experience.

The early deaf dataset, acquired using a multi-echo fMRI protocol, was preprocessed using fMRIPrep (Esteban et al., 2019) in combination with TEDANA (DuPre et al., 2021). This workflow enables echo combination and TE-dependent ICA denoising, improving BOLD sensitivity and effectively removing non-BOLD artifacts specific to multi-echo acquisitions.

For both datasets, preprocessing included anatomical preprocessing, head-motion correction, anatomical–functional coregistration, projection of functional time series onto the cortical surface (HCP fsLR_32k), nuisance regression, temporal filtering, and surface-based spatial smoothing. All dataset-specific preprocessing steps, parameter choices, and masking procedures are described in detail in the Supplementary materials.

### Sensory integration model

We applied the sensory integration model described in (Wei et al., 2024), which is grounded in the anchoring role of primary sensory cortex within the sensory processing organization. To quantify sensory integration, we used a generalized linear model (GLM) with non-negative constraints, employing the averaged time series from the primary visual cortex (V1), primary somatosensory cortex (S1), and primary auditory cortex (A1) as predictors for the time series at each cortical surface vertex. To evaluate potential multicollinearity among these primary sensory signals, Variance Inflation Factors (VIF) were calculated. All VIFs were substantially below the conventional threshold for collinearity (VIF = 5), indicating minimal multicollinearity. Full results are reported in Supplementary Table 1 and Supplementary Fig. 2. Primary sensory areas were defined using Glasser’s multimodal parcellation (Glasser et al., 2016): parcel V1 corresponds to the primary visual cortex, parcel A1 to the primary auditory cortex, and parcels 1, 2, 3a, and 3b to the primary somatosensory cortex.

The GLM is represented by the following equation,

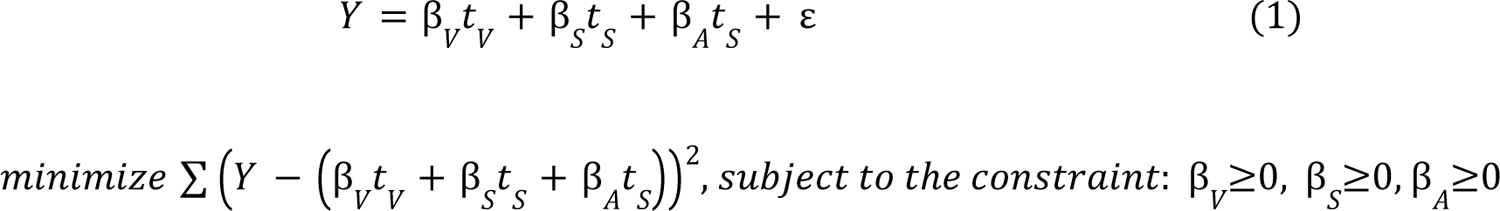

*Y is the time series of the vertex. t_V_, t_S_, and t_A_ are averaged time series of V1, S1, and A1, separately. β_V_, β_S_, and β_A_ are the non-negative regression coefficients.* ε *is the noise, assumed to be Gaussian-distributed with mean 0, it represents the variance not explained by the primary sensory signal*.

The regression coefficients (β_V_, β_S_ and β_A_) obtained from Equation 1 represent the contribution of each specific sensory modality at each cortical vertex. We refer to these coefficients collectively as “sensory parameters.” The two dimensions of our sensory integration model were derived from these sensory parameters.

Given that the early blind datasets were collected across different research sites, exhibiting age and sex variability among subjects, we included age and sex as additional covariates when estimating sensory parameters for the early blind datasets. In contrast, age and sex are matched across groups in the early deaf dataset and therefore were not included as covariates.

As the first dimension of the sensory integration model, sensory magnitude reflects the sensory processing hierarchy based on the proportion of variance explained by the primary sensory signals.

To reduce sensitivity to the non-uniform distribution of explained variance and to emphasize relative differences across cortical regions, these values were ranked and rescaled to a standardized range between 0 and 1 to define sensory magnitude.

The second dimension, sensory angle, characterizes the relationship between different sensory modalities by converting the sensory parameters into angular measures. This transformation positions the primary sensory areas in a “sensory-defined color space” capturing transformations from “purely visual”, “purely auditory” and “purely somatosensory” to multisensory regions and then to fully transmodal. The equations used to transform the three sensory parameters into angles are provided below.

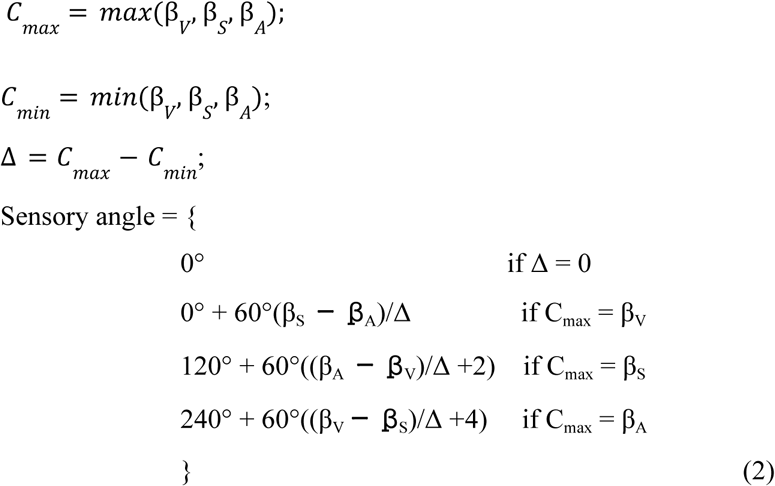

*C_max_ is the maximum sensory parameter. C_min_ is the minimum sensory parameter. Δ is the difference between C_max_ and C_min_. Sensory angle is the angular position in a unit circle with a range from 0° to 360°*.

To investigate the network-level consequences of altered sensory integration, we performed a seed-based functional connectivity analysis. The regions of interest (ROIs) were defined as the clusters exhibiting significant between-group differences in sensory integration metrics (sensory angle or magnitude) identified in the previous step. For each subject, we computed the functional connectivity maps originating from these specific ROIs to determine how local deviations in sensory processing extend to large-scale network organization.

### Functional gradients

Functional gradients, derived from decomposing whole-brain functional connectivity profiles, have been demonstrated as effective measures capturing principal axes of functional organization (Margulies et al., 2016). To derive a consensus map of functional organization across early sensory deprived individuals and controls, we applied Generalized Canonical Correlation Analysis (GCCA) (Afshin-Pour et al., 2012) via the mvlearn package (Perry et al., 2020) to calculate functional gradients. A significant challenge in gradient analysis is the ‘correspondence problem,’ where gradients derived independently for each subject may be rotated or ordered differently, precluding direct comparison. GCCA overcomes this by identifying a common latent manifold while simultaneously calculating subject-specific projections into that shared space. This unified coordinate system allows for vertex-wise statistical testing of individual gradient maps across groups without the need for post-hoc Procrustes rotation or other alignments. We focused our analysis on the first three gradients, as these components capture the strongest shared functional features between early sensory deprived individuals and controls (Supplementary Fig. 3 and 4)

Both the sensory integration model and functional gradients characterize alterations in functional organization associated with sensory deprivation. To assess the correspondence between these complementary approaches, we tested the spatial similarity of results derived from the sensory integration model, functional connectivity, and functional gradients. However, since the spatial extent of the effects identified by each metric varies significantly, a standard overlap measure could be biased. Therefore, we applied an inverse size weighting scheme to the Dice coefficient, ensuring that differences in cluster size do not disproportionately penalize the similarity score. The formula for calculating the modified coefficient is described below:

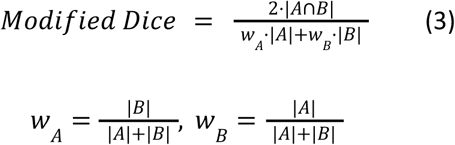

Where, |A| and |B| are the sizes of the results obtained from different metrics, w_A_ and w_B_ are the weights assigned to result A and result B, respectively.

### Statistical comparison

To compare the global topography of sensory integration metrics, we assessed the spatial similarity of group-level maps. For Sensory Angle, we utilized circular correlation (Jammalamadaka & Sengupta, 2001), while for Sensory Magnitude, we employed Spearman rank correlation. Group-level Sensory Angle maps were derived by averaging individual angles. For Sensory Magnitude, we calculated the mean proportion of variance explained by primary sensory signals across participants. To capture relative positioning along the cortical hierarchy while minimizing sensitivity to distributional skew and outliers, we ranked these values, and rescaled them to a range of 0 to 1 to obtain standardized group-level magnitudes.

At the vertex level, we examined group differences using statistical models tailored to the data type and cohort structure. For the early blind dataset, we employed a 3 (Site: Montreal, Trento, Paris) × 2 (Group: Early Blind, Sighted Control) two-way between-subjects ANOVA. Given the circular nature of Sensory Angle, we used the Harrison–Kanji test (Harrison & Kanji, 1988) as the circular equivalent of the two-way ANOVA. For Sensory Magnitude, a standard linear two-way ANOVA was applied. For the early deaf dataset, comparisons across the three participant groups were conducted using a one-way design: the Watson–Williams test (Watson & Williams, 1956) was used for Sensory Angle, while a standard one-way ANOVA was used for Sensory Magnitude. Functional connectivity and functional gradient maps were analyzed using the standard linear ANOVA models described above.

To control for demographic variability in the multisite early blind dataset, age and sex were included as covariates in these statistical models for comparing functional connectivity and gradient values across groups. Crucially, age and sex were not included as covariates in the statistical comparisons of Sensory Angle and Magnitude. This exclusion was intentional, as these variables were already included as predictors in the voxel-wise linear models used to estimate the sensory parameters, thereby removing their confounding effects prior to group-level analysis. Notably, age and sex were matched across groups in the early blind dataset and therefore were not included as covariates for the statistical comparisons.

To identify brain regions exhibiting significant group differences, we employed a cluster-based permutation test to correct for multiple comparisons. We generated a null distribution of cluster sizes by performing 5,000 permutations in which group labels were randomly shuffled and the statistical maps recomputed. For each iteration, maps were thresholded at a vertex-wise p < 0.001, and the maximum suprathreshold cluster size was recorded. Significance was determined by the 95th percentile of this distribution, and only observed clusters exceeding this critical extent were retained.

## Results

To investigate the impact of early sensory deprivation on the functional organization of the cortex, we analyzed resting-state fMRI data from two independent cohorts: individuals with early blindness and individuals with early deafness. We employed a multimodal framework combining our function-based Sensory Integration Model with functional gradient mapping. This approach allows us to simultaneously characterize alterations in sensory integration and the resilience of the brain’s intrinsic functional topology.

The Sensory Integration Model quantifies the functional role of cortical areas through two dimensions: Sensory Angle, which reflects the proportional contribution of visual, auditory, and somatosensory modalities, and Sensory Magnitude, which indexes the hierarchical progression from sensory dependence to abstract integration. Complementing this, we applied Generalized Canonical Component Analysis (GCCA) to derive functional gradients that represent the brain’s dominant functional features. By integrating these metrics, we examined both the global spatial similarity and vertex-wise differences between deprived individuals and controls, aiming to uncover how the loss of a sensory modality reshapes the interplay between sensory integration strategies and the cortical functional architecture.

### Early blindness

#### Sensory integration mapping

The sensory integration mapping was visualized in polar coordinate space (Fig. 1a) and on the cortical surface (Fig. 1b). Qualitative inspection of the plots indicated that early blind individuals reduced angular dispersion, with vertices clustering more tightly around the visual anchoring angle (0°) compared to sighted controls. This indicated a reduction in the angular dispersion of the sensory profile, confirming that the primary distinction between groups involved a shift toward visual-dominant processing. This shift is likely to reflect increased functional processing of information from V1 rather than processing of visual input, which is absent in the blind. Commensurate with this visual shift, the areas surrounding V1 exhibited a more reddish hue in the early blind compared to sighted controls (Fig. 1b), reflecting a greater similarity to V1 during the resting state.

**Figure 1.**
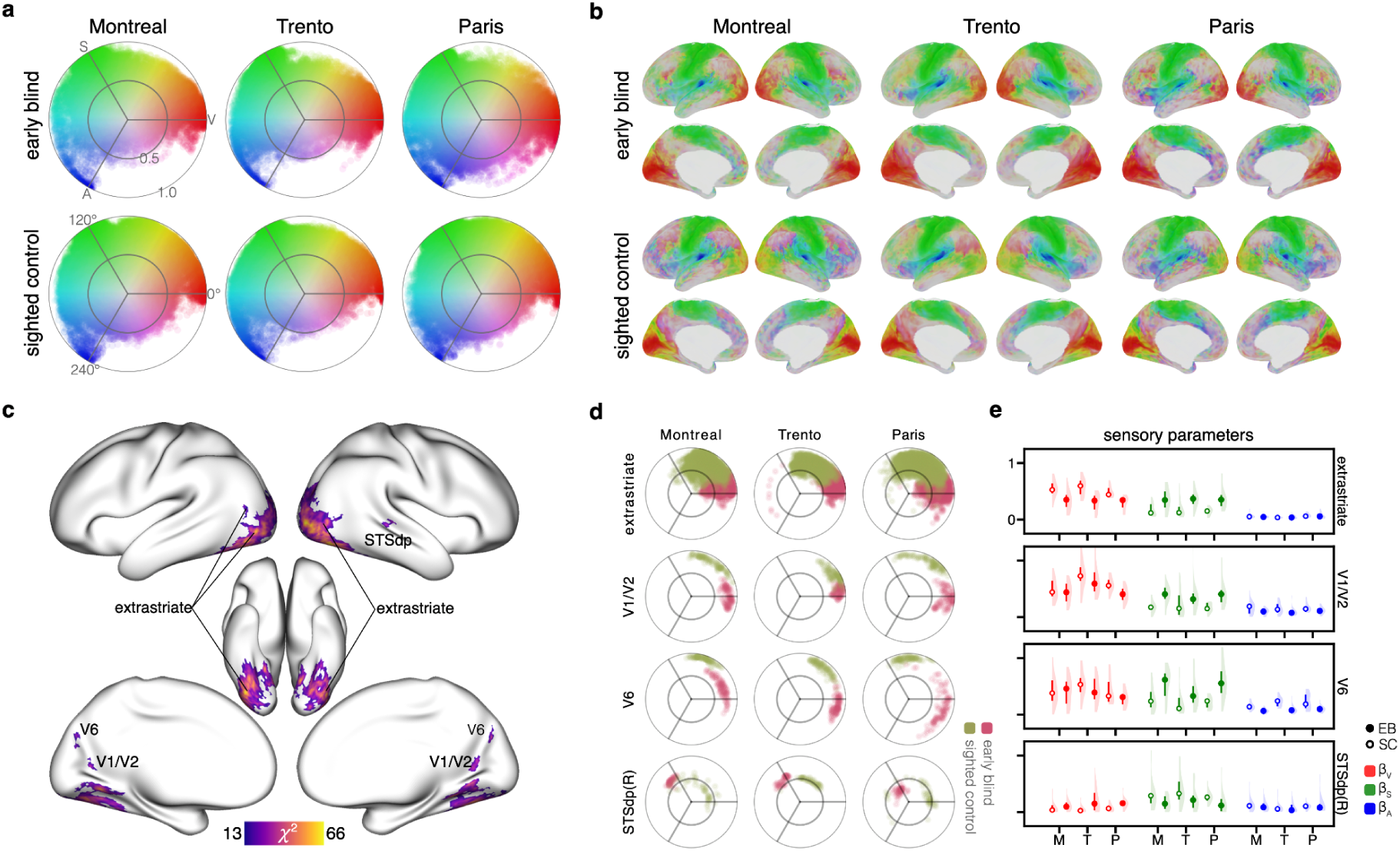
Sensory integration mapping and cross-modal plasticity in early blind and sighted individuals. **a)** Sensory integration mapping in polar space. Colors are defined by an HSV model: hue represents sensory angle, saturation represents sensory magnitude, and brightness is constant at 0.86. b) Group-level sensory integration maps projected onto the cortical surface. c) Cortical regions showing significant differences in sensory angle between early blind and sighted controls. d) Cluster-based sensory integration patterns. Polar plots show the distribution of the group-level sensory angles and magnitudes within each cluster, each point representing a vertex. e) Cluster-based sensory parameter distributions of sensory parameters across clusters and study sites. Point-interval plots show the median (points) and interquartile range (vertical lines) for each group, overlaid on shaded kernel density estimates (shadows) representing the distribution of subject-wise beta weights. Early blind subjects are indicated by open circles and sighted controls by solid circles. STSdp, dorsolateral superior temporal sulcus; M, Montreal dataset; T, Trento dataset; P, Paris dataset; EB, early blind; SC, sighted control; β_V_, visual parameter; β_S_, somatosensory parameter; β_A_, auditory parameter.

To compare the global topology of sensory integration across groups and datasets, we quantified cross-site similarities of group-level Sensory Angle and Sensory Magnitude maps across the three independent early blind datasets (Table 2). Sensory Angle maps showed moderate but structured cross-site similarities, with consistent high and stable values within the early blind group (EB–EB: r ≈ 0.69–0.72) and within sighted groups (SC–SC: r ≈ 0.64–0.67). Between-group similarities (EB–SC: r ≈ 0.61–0.70) fell within a comparable range as within-groups similarities but exhibited greater variability across site combinations. This pattern indicates that early blindness is associated with a stable and site-independent alteration in sensory integration profiles. In contrast, Sensory Magnitude maps were highly similar across all datasets and groups (r ≈ 0.87–0.94). Together with sensory angle findings, it suggested that early sensory deprivation primarily alters the relative weighting of sensory modalities rather than the overall magnitude of sensory integration. In other words, despite alterations in sensory integration within early blind individuals, the typical hierarchical organization of sensory processing, indexed by the consistent magnitude, was largely preserved at a global topographic level.

**Table 2.** Group-level comparison of sensory angles and magnitudes for early blind datasets.

| **panel a. Similarity of group-level sensory angles** |  |  |  |  |  |  |  |
| --- | --- | --- | --- | --- | --- | --- | --- |
|  |  | Montreal |  | Trento |  | Paris |  |
|  |  | EB | SC | EB | SC | EB | SC |
| Montreal | EB | 1 |  |  |  |  |  |

|  |  |  |  |  |  |  |  |
| --- | --- | --- | --- | --- | --- | --- | --- |
|  | SC | 0.635 | 1 |  |  |  |  |
| Trento | EB | 0.722 | 0.637 | 1 |  |  |  |
|  | SC | 0.65 | 0.673 | 0.709 | 1 |  |  |
| Paris | EB | 0.686 | 0.63 | 0.722 | 0.677 | 1 |  |
|  | SC | 0.558 | 0.636 | 0.611 | 0.652 | 0.625 | 1 |
| **panel b. Similarity of group-level sensory magnitudes** |  |  |  |  |  |  |  |
|  |  | Montreal |  | Trento |  | Paris |  |
|  |  | EB | SC | EB | SC | EB | SC |
| Montreal | EB | 1 |  |  |  |  |  |
|  | SC | 0.907 | 1 |  |  |  |  |
| Trento | EB | 0.892 | 0.898 | 1 |  |  |  |
|  | SC | 0.902 | 0.905 | 0.94 | 1 |  |  |
| Paris | EB | 0.875 | 0.865 | 0.888 | 0.904 | 1 |  |
|  | SC | 0.884 | 0.902 | 0.906 | 0.917 | 0.894 | 1 |

*EB, early blind; SC, sighted control.*

Vertex-wise comparisons were performed to ascertain regions exhibiting significant differences in sensory integration between the early blind and sighted controls. No significant between-group differences were identified in Sensory Magnitude, further supporting the preservation of the processing hierarchy. Conversely, significant differences in Sensory Angles were observed in bilateral extrastriate cortex, including multiple dorsal and lateral visual subregions (V2, V3, V4, V8, VMV, VVC, FFC, PIT, PH, LO1–LO3, and MT/MST), as well as in V1/V2, V6, and the right dorsal posterior superior temporal sulcus (STSdp) (Fig. 1c and Table 3).

**Table 3.** Cluster-based sensory angular changes.

|  | **Montreal** |  | **Trento** |  | **Paris** |  | **cluster size** | **p-value** |
| --- | --- | --- | --- | --- | --- | --- | --- | --- |
|  | mean angle |  | mean angle |  | mean angle |  |  |  |
|  | EB | SC | EB | SC | EB | SC |  |  |
| extrastriate (L) | 16.4° | 59.1° | 19.4° | 68.4° | 6.8° | 68.5° | 1524 | p < 0.0002 |
| extrastriate (R) | 14.1° | 65.1° | 15.2° | 68.3° | 12° | 68° | 1648 | p < 0.0002 |
| V1/V2 (L) | 2.6° | 62.3° | 9.3° | 32° | 344.3° | 51.3° | 37 | 0.032 |
| V1/V2 (R) | 359.4° | 58° | 6.5° | 32.9° | 4.1° | 67.1° | 64 | 0.0092 |
| V6 (L) | 23.3° | 65.4° | 356.2° | 48.1° | 338.6° | 88.4° | 50 | 0.0172 |
| V6 (R) | 18.5° | 80.7° | 342.2° | 61.4° | 355.8° | 96.4° | 43 | 0.0234 |
| STSdp (R) | 134.7° | 54.9° | 128.6° | 63° | 129.2° | 20.7° | 52 | 0.0144 |

L, left hemisphere; R, right hemisphere; EB, early blind; SC, sighted control.

The subsequent analysis of the cluster-level patterns (Fig. 1d) characterized the precise nature of these angular shifts. For the significant clusters located within the visual cortex (extrastriate, V1/V2, and V6), the angular patterns of early blind individuals shifted away from the predominantly visual-somatosensory angular range observed in controls toward the visual-dominant range (near 0°). Conversely, the pattern of the STSdp cluster shifted toward the somatosensory-dominant range (near 120°). This counter-intuitive finding—a strong shift toward visual dominance despite lifelong visual deprivation—indicates that the synchronization of intrinsic brain activity between V1 and the extrastriate cortex was higher in early blind individuals than in the sighted controls.

To further dissect the factors driving these Sensory Angle alterations, we directly compared the sensory parameters (the beta values derived from the GLM) within each significant cluster (Fig. 1e and Supp. Tab. 2). The pattern shift in the extrastriate cluster was primarily driven by a significantly increased visual parameter and a corresponding decreased somatosensory parameter. In contrast, the pattern shifts of the V1/V2 and V6 clusters were driven primarily by the decreased somatosensory parameters. In the case of the STSdp cluster, the shift was driven by a slight decrease in the visual parameter and a robust increase in the somatosensory parameter. These data demonstrate that the sensory angle differences are mediated by a complex rebalancing: the increased contribution from the visual modality and the decreased contribution from the somatosensory modality concurrently lead to the observed reorganization in the early blind visual and temporoparietal cortex.

### Functional connectivity profiles of regions with altered sensory integration

To investigate the network-level consequences of altered sensory integration, we next investigated functional connectivity from the clusters with significant sensory angle differences between groups.

Functional connectivity analyses seeded from the extrastriate cluster revealed widespread differences between early blind and sighted controls. Those differences were localized to V1, V3/V6, parahippocampal gyrus, somatosensory/motor regions, superior temporal gyrus, middle temporal area, inferior frontal gyrus, i6-8/s6-8, anterior ventral insula, inferior parietal sulcus, posterior inferior temporal gyrus, dorsal medial prefrontal cortex, and posterior cingulate cortex (Fig. 2a). More specifically, compared with sighted controls, the early blind group showed stronger functional connectivity between extrastriate cortex and higher-order regions within the FPN and DMN, alongside weaker functional connectivity between extrastriate cortex and sensorimotor-related regions (Fig. 2b). For the other two visual-related clusters, the V1/V2 cluster of early blind individuals exhibited lower connectivity with bilateral somatosensory/motor regions and right A5 (Fig. 2c). The V6 cluster demonstrated lower connectivity to somatosensory/motor regions as well as visual and higher-order regions, except for the left V2/V3/V4 and right anterior ventral insula (AVI) (Fig. 2d). For the STSdp cluster, the early blind individuals exhibited lower connectivity to visual and sensorimotor regions and a higher connectivity to the parietal-occipital sulcus (Fig. 2e). Overall, clusters exhibiting significantly altered sensory angles in the early blind demonstrated enhanced connectivity with higher-order cortical regions, and reduced connectivity with somatosensory/motor areas during the resting state.

**Figure 2.**
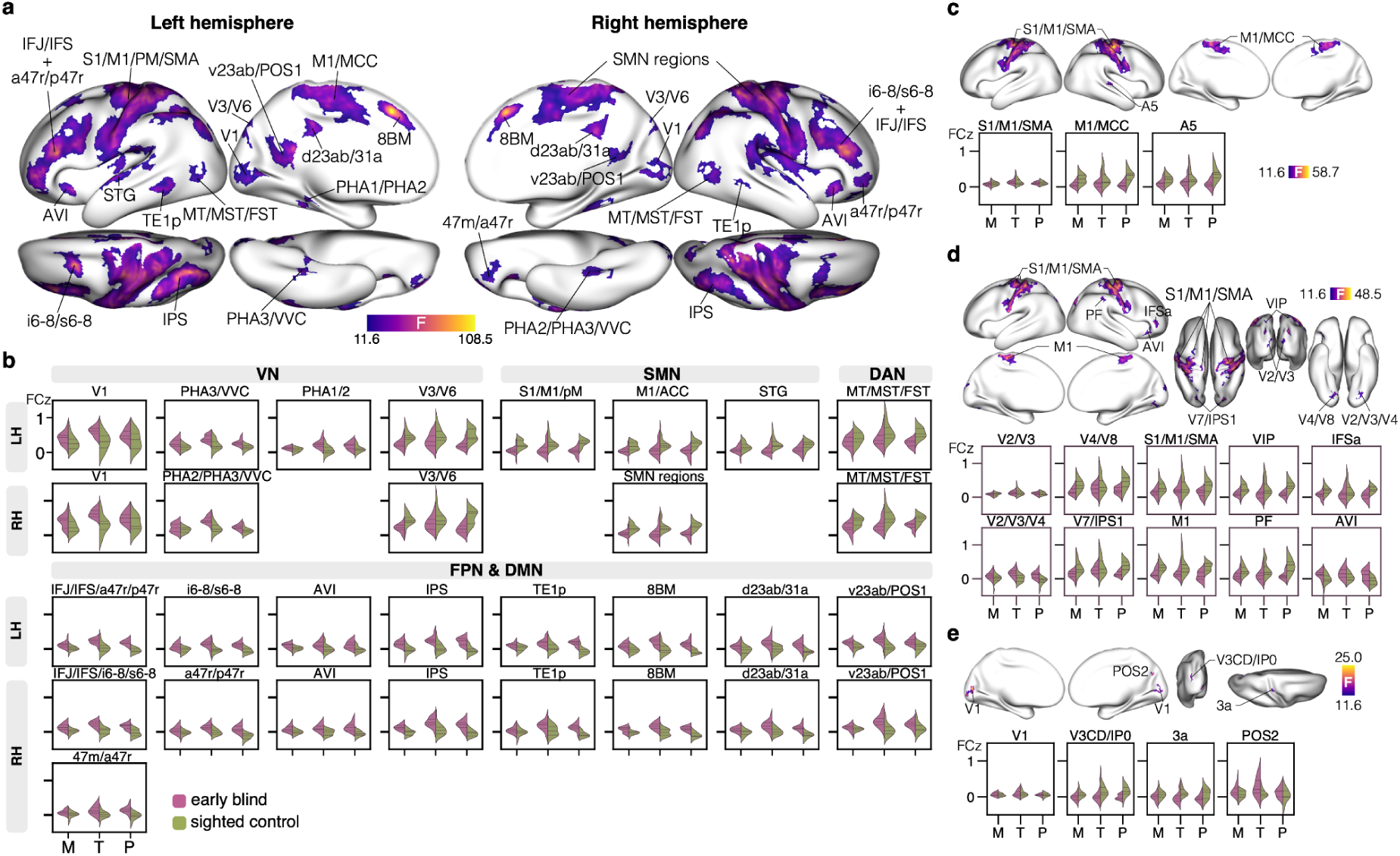
Significant differences of the cluster-based functional connectivity between early blind and sighted controls. **a)** Significant differences of the extrastriate-based FC. b) Distribution of the extrastriate-based FC within each significant cluster. The shaded area of each violin plot represents the data density, while horizontal lines inside each violin indicate the 25th, 50th (median), and 75th percentiles. c) Significant differences of the V1/V2-based FC. d) Significant differences of the V6-based FC. e) Significant differences of the STSdp-based FC. VN, visual network; SMN, sensorimotor network; DAN, dorsal attention network; FPN, frontoparietal network; DMN, default mode network; M, Montreal dataset; T, Trento dataset; P, Paris dataset; LH, left hemisphere; RH, right hemisphere.

### Functional gradient profiles

To derive a consensus map of functional organization, we employed Generalized Canonical Component Analysis (GCCA) to calculate functional gradients. This method aligns individual data into a common latent space to identify spatial patterns that are highly consistent across all participants. We focused our analysis on the first three gradients, as these components capture the strongest shared functional features across early blind individuals and sighted controls (Supp. Fig. 3). The first gradient (G1) represents the principal hierarchy extending from unimodal sensory-motor regions to transmodal association cortex. Significant differences between early blind and sighted controls in G1 were observed primarily in the bilateral extrastriate cortex and left area 6a, which were located at the primary end of the hierarchy, excluding V1 as the sensory angle result. The second gradient (G2) captures the sensory dissociation between visual and somatomotor processing streams. The significant G2 differences were identified in the visual cortex, encompassing V1 and extrastriate regions, as well as somatosensory regions, which were located at both ends of G2 distribution. Collectively, G1 and G2 mainly showed differences in the extrastriate cortex. In contrast, G3 exhibited a broader and more distributed pattern involving sensory, motor, and higher-order association regions (Fig. 3a). Importantly, after identifying an extrastriate cluster in the sensory angle comparison (Fig. 1c), we compared its functional connectivity maps (Fig. 2a). The resulting difference map closely resembled the spatial distribution of significant group differences observed in G3 (Fig. 3a). This correspondence with the sensory integration results supports the robustness of these findings.

**Figure 3.**
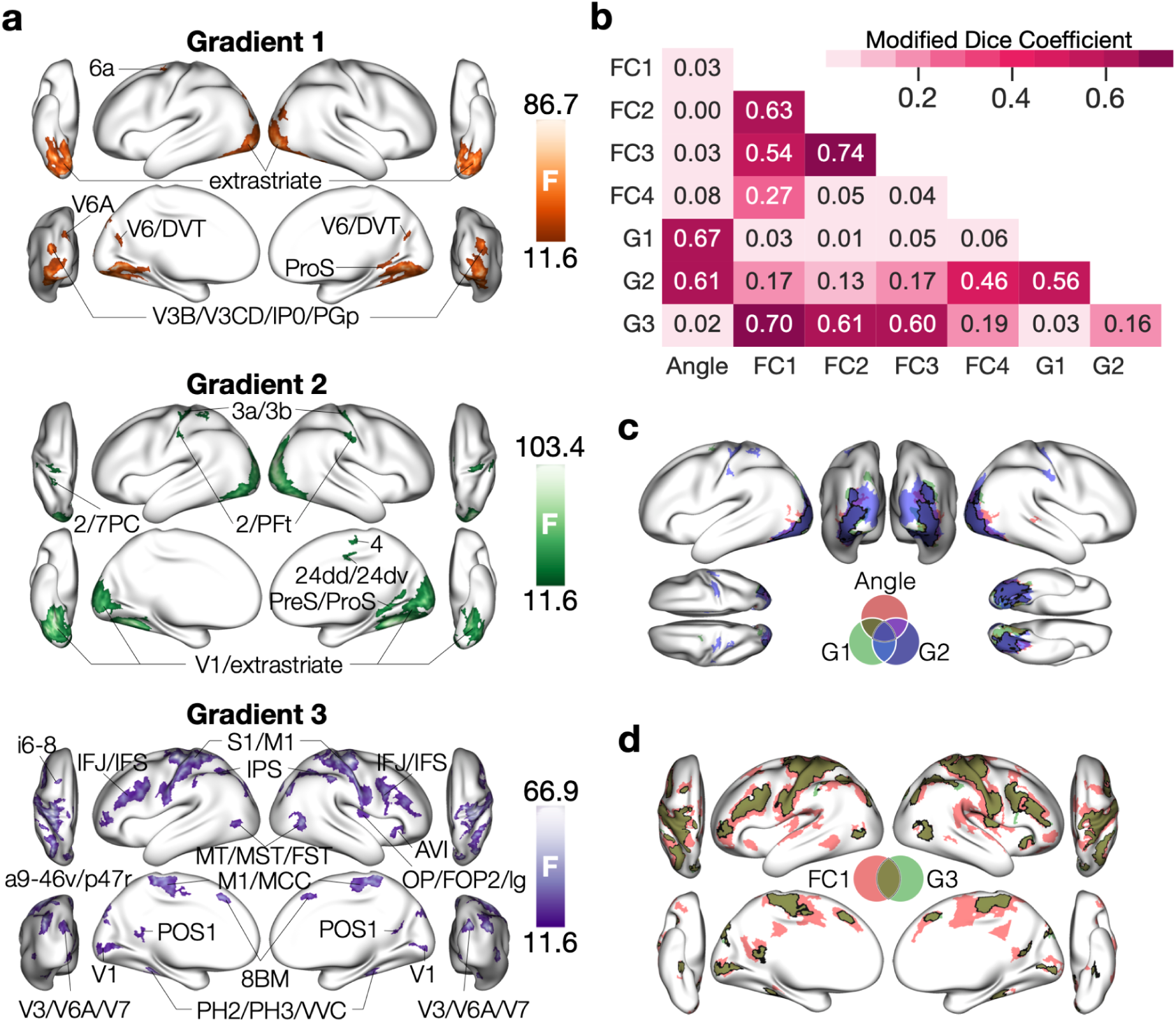
Significant differences of functional gradients between early blind and sighted controls and its overlap with other results. **a)** Significant differences of functional gradients on the cortical surface. b) Spatial overlap ratio between sensory angle, functional connectivity, and functional gradient results. c) Spatial overlap between sensory angle, Gradient 1, and Gradient 2 results. d) Spatial overlap between the extrastriate-based FC and Gradient 3 results. G1, the principal gradient, G2, the secondary gradient; G3, the third gradient; FC1, the extrastriate-based connectivity; FC2, V1/V2-based connectivity; FC3, V6-based connectivity; FC4, STSdp-based connectivity.

### Spatial correspondence between sensory integration and functional gradient results

To further assess the consistency between the macroscale functional features (Gradients) and intrinsic sensory processing strategies (Sensory Integration Model), we quantified the spatial overlap of their significant group differences. Given the variation in the spatial extent of the effects identified by each metric, we employed an inverse size-weighted Dice coefficient to ensure a balanced comparison.

First, we examined the gradients representing the primary axes of sensory organization. We found that the reorganization patterns of Gradient 1 (G1) and Gradient 2 (G2) exhibited the highest spatial overlap with the Sensory Angle results (Fig. 3c). Surface visualization confirmed that these overlapping alterations were predominantly localized to the extrastriate cortex (Fig. 3d). This suggests that the shifts in intrinsic sensory dominance (Sensory Angle) are spatially congruent with alterations in the brain’s fundamental sensory processing axes.

Next, we examined the higher-order Gradient 3 (G3). Here, the highest overlap was observed with the extrastriate-based functional connectivity (FC) patterns. While connectivity maps seeded from V1/V2 and V6 also showed partial overlap with G3, specifically within the primary somatosensory cortex (S1), the extrastriate-based connectivity demonstrated a far more extensive spatial correspondence. Crucially, this overlap extended beyond S1 to include distributed regions such as the anterior ventral insula (AVI), aligning closely with the topography of G3 (Fig. 3e).

Collectively, these patterns demonstrate a high degree of spatial correspondence between the alterations in sensory integration related metrics (Angle, Connectivity) and the reorganization of the brain’s dominant functional features.

### Early deafness

#### Sensory integration mapping

The sensory integration mapping was visualized in polar coordinate space and on the cortical surface (Fig. 4a). Qualitative inspection indicated that early deaf individuals exhibited a tighter clustering of vertices around the auditory anchoring angle (240°) compared to hearing controls. This suggests that the primary functional distinction involves the tuning of auditory processing circuits. Commensurate with this shift, the superior temporal area adjacent to primary auditory cortex (A1) exhibited a magenta hue in early deaf individuals which is distinct from the green hue observed in hearing controls (Fig. 4a), indicating intrinsic brain activity combining information from auditory and visual areas.

**Figure 4.**
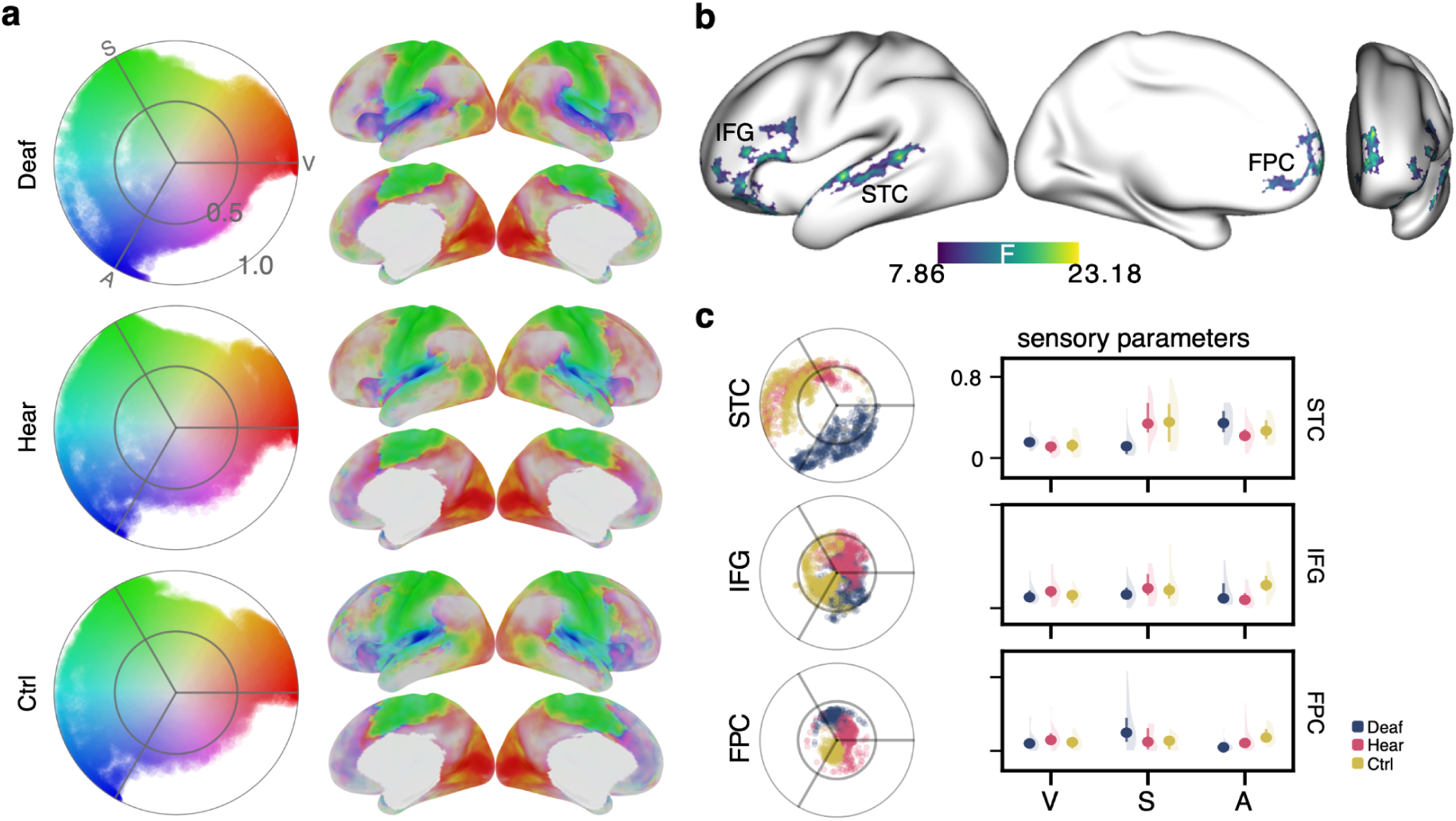
Sensory integration mapping and its between-groups differences. **a)** Sensory integration mapping visualized in polar space and projected onto the cortical surface. b) Cortical regions showing significant sensory angle differences among the three groups of the early deaf dataset. c) Cluster-based sensory integration patterns and sensory parameter distributions. Left: Polar plots showed the distributions of group-level sensory angles and magnitudes within each cluster, each point representing a vertex. Right: Distributional profiles of sensory parameters. Point-interval plots show the group-level median (points) and interquartile range (vertical lines), overlaid on kernel density estimates (shadows) representing subject-wise beta weights for visual, somatosensory, and auditory modalities. Deaf, early deaf with knowledge of cued speech; Hear, hearing controls with knowledge of cues speech; Ctrl, hearing controls without knowledge of cued speech; STC, superior temporal cortex; IFG, inferior frontal gyrus; FPC, frontopolar cortex; V, visual sensory parameter; S, somatosensory sensory parameter; A, auditory sensory parameter.

The early deaf dataset comprised three distinct groups: early deaf individuals with knowledge of cued speech (Deaf), hearing controls with knowledge of cued speech (Hear), and hearing controls without such knowledge (Ctrl). This design allowed us to disentangle functional reorganization driven by sensory deprivation from that driven by language experience.

To quantitatively compare the global topology of sensory integration, we computed between-group similarities of the group-level Sensory Angle and Magnitude maps (Tab. 4). Consistent with the early blind analysis, the spatial similarity of Sensory Magnitude was consistently high across all pair-wise comparisons. However, the similarity of Sensory Angles was higher when comparing the two hearing groups (Hear vs. Ctrl) than when comparing the deaf group to either hearing group. This finding reinforces the conclusion that while sensory integration strategies (Angle) are susceptible to both deprivation and language experience, the fundamental hierarchical organization (Magnitude) remains largely preserved.

**Table 4.** Global-level comparison.

| **Table 4. Global-level comparison** |  |  |  |
| --- | --- | --- | --- |
| **panel a. sensory angle correlation** |  |  |  |
|  | Deaf | Hear | Ctrl |
| Deaf | 1 |  |  |
| HC with CS | 0.663 | 1 |  |
| HC without CS | 0.588 | 0.726 | 1 |
| **panel b. sensory magnitude correlation** |  |  |  |
|  | Deaf | Hear | Ctrl |

| Table 4. Global-level comparison |  |  |  |
| --- | --- | --- | --- |
| Deaf | 1 |  |  |
| HC with CS | 0.957 | 1 |  |
| HC without CS | 0.946 | 0.959 | 1 |

CS, cued speech.

Vertex-wise comparisons were performed to identify regions showing significant group differences. No significant differences were found in Sensory Magnitude. Significant differences in Sensory Angles were localized to the left superior temporal cortex (STC), left inferior frontal gyrus (IFG), and left frontopolar cortex (FPC) (Fig. 4b).

Post-hoc comparisons revealed a functional dissociation between these clusters, attributing them to distinct plastic mechanisms (Tab. 5). The STC cluster exhibited significant differences between early deaf individuals (Deaf) and both hearing control groups (Hear and Ctrl), regardless of language experience, indicating a specific effect of sensory deprivation. In contrast, the IFG and FPC clusters were associated with differences between Cued Speech users (Deaf and Hear) versus non-users (Ctrl), highlighting the influence of language experience rather than sensory loss. This dissociation demonstrates that sensory integration mapping is sensitive not only to reorganization driven by sensory deprivation but also to plasticity induced by specific language modalities.

**Table 5.** Post-hoc comparison of sensory angles within each significant cluster.

|  | Deaf vs Hear vs Ctrl |  | Deaf vs Hear |  | Deaf vs Ctrl |  | Hear vs Ctrl |  |
| --- | --- | --- | --- | --- | --- | --- | --- | --- |
|  | F | p | F | p | F | p | F | p |
| STC | 21.704 | 1.12e-07 | 47.273 | 4.16e-08* | 24.145 | 2.1e-05* | 0.327 | 0.57 |
| IFG | 18.516 | 7.11e-07 | 2.38 | 0.13 | 15.5 | 3.74e-04* | 36.48 | 5.02e-07* |
| FPC | 19.467 | 4.05e-07 | 4.131 | 0.05 | 39.98 | 2.9e-07* | 16.848 | 2.07e-04* |

*Deaf, early deaf with knowledge of cued speech; Hear, hearing controls with knowledge of cues speech; Ctrl, hearing controls without knowledge of cued speech; STC, superior temporal cortex; IFG, inferior frontal gyrus; FPC, frontopolar cortex. \* $p < 0.0167$ (Bonferroni correction for comparisons of the averaged sensory angles within each cluster, $0.05/3 = 0.0167$ )*

Subsequently, we analyzed the cluster-level patterns to characterize the specific nature of these angular shifts (Fig. 4c). In the STC, which reflected the specific effect of sensory deprivation, the angular profile of the early deaf group (Deaf) was predominantly situated within the auditory-visual angular range (mean angle = 277.5°). In distinct contrast, the patterns of the two hearing groups were positioned within the auditory-somatosensory angular range (Hear: 137.9°; Ctrl: 158.8°).

Regarding the clusters associated with language experience, the IFG and FPC exhibited distinct profiles based on the knowledge of Cued Speech rather than hearing status. For the IFG, the group without Cued Speech knowledge (Ctrl) was situated in the somatosensory-auditory range (218.2°). Conversely, the groups with Cued Speech knowledge shifted markedly toward the visual anchoring angle: the early deaf group (Deaf) was located in the visual-dominant range (354°) and the hearing Cued Speech group (Hear) in the visual-somatosensory range (48.1°). Similarly, in the FPC, the non-Cued group (Ctrl) was situated in the auditory-dominant range (251.7°), whereas the Cued Speech groups exhibited distinct profiles situated in somatosensory-(Deaf: 85.7°) or visual-related (Hear: 359.4°) ranges.

Collectively, these findings reveal a convergent trajectory of reorganization. The patterns associated with deafness (in the STC) shifted from the somatosensory-auditory range observed in controls toward the auditory-visual range (240°–360°). Concurrently, the patterns reflecting Cued Speech experience (in the IFG) shifted from the somatosensory-auditory range toward the visual anchoring angle (0°). These results confirm that the observed functional changes in early deaf individuals are characterized by the specific recruitment of visual processing within the auditory and language networks, supporting both cross-modal plasticity and the demands of a visual-gestural language system.

### Functional connectivity and gradient profiles

To elucidate the network-level consequences of the observed local reorganization, we evaluated the functional connectivity profiles using the significant clusters identified by Sensory Angle analysis (STC, IFG, and FPC) as seeds (Fig. 5). The STC-based connectivity exhibited significant between-group differences located within the bilateral somatosensory/motor cortex, right V4t/FST, and right A4. For the clusters associated with language effects, the IFG seed in early deaf individuals showed reduced connectivity to the left primary somatosensory cortex (S1), while the FPC seed showed reduced connectivity to the bilateral superior temporal gyrus (STG).

**Figure 5.**
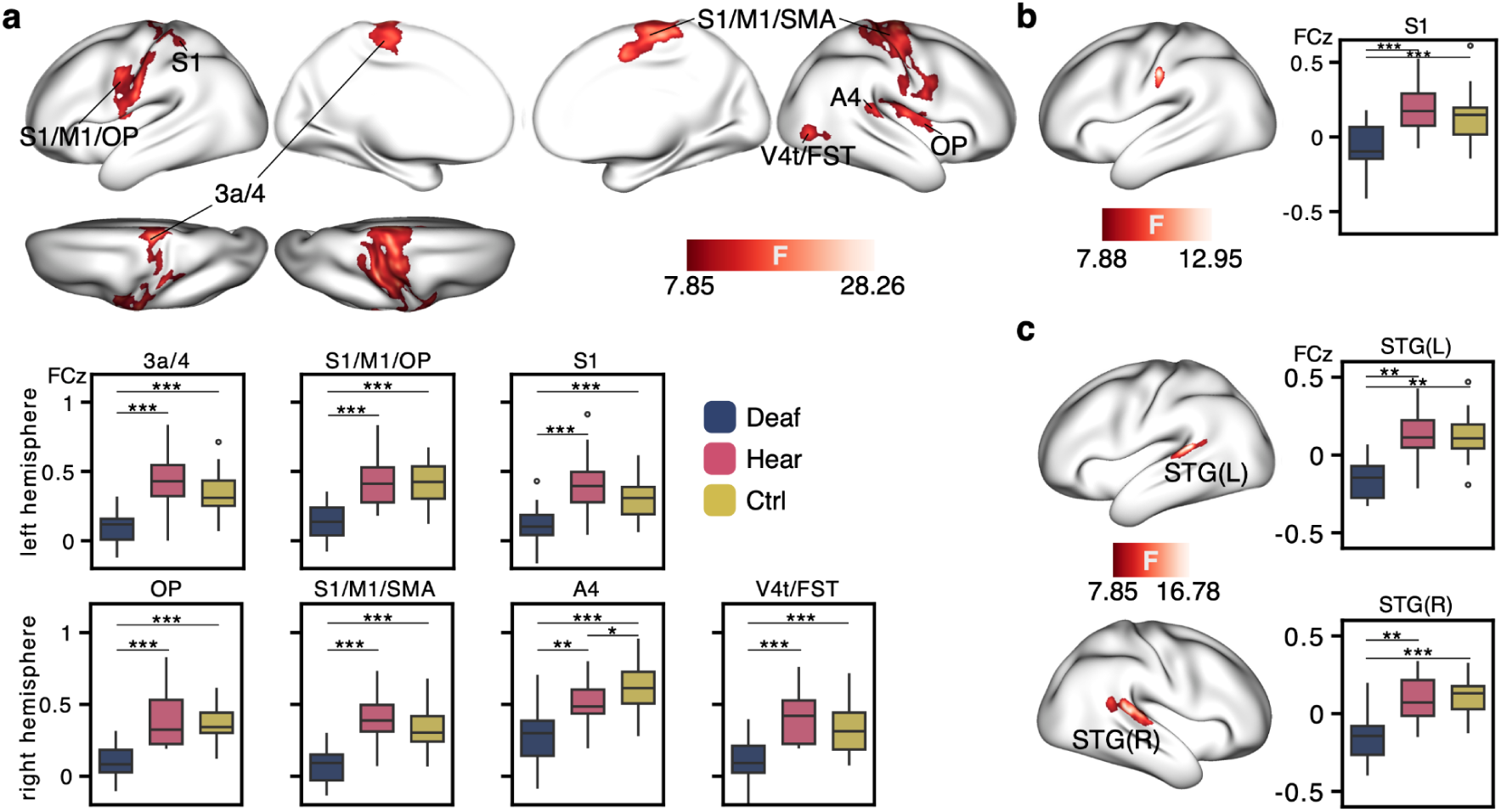
Significant between-group differences of the cluster-based functional connectivity. **a)** Significant between-group differences of the STC-based FC. b) Significant between-group differences of the IFG-based FC. c) Significant between-group differences of the FPC-based FC. Deaf, early deaf with knowledge of cued speech; Hear, hearing controls with knowledge of cues speech; Ctrl, hearing controls without knowledge of cued speech.

Crucially, post-hoc analysis revealed that all significant between-group differences in functional connectivity were driven exclusively by the effect of sensory deprivation (differences between early deaf individuals and both hearing groups). This held true even for the IFG and FPC seeds, which had originally been identified based on language-related differences in the Sensory Angle analysis. Furthermore, whereas early blindness was associated with specific enhancements in connectivity, early deaf individuals exhibited a generalized pattern of reduced connectivity relative to controls, suggesting a distinct mechanism of macroscopic reorganization.

We subsequently compared the first three GCCA-based functional gradients, which captured the highest shared structure strength across the cohort (Supplementary Fig. 4). No significant between-group differences were found in Gradient 1. However, significant differences in Gradient 2 and Gradient 3 were identified in the bilateral superior temporal gyrus, which spatially overlapped with the Sensory Angle results in the left hemisphere (Fig. 6).

**Figure 6.**
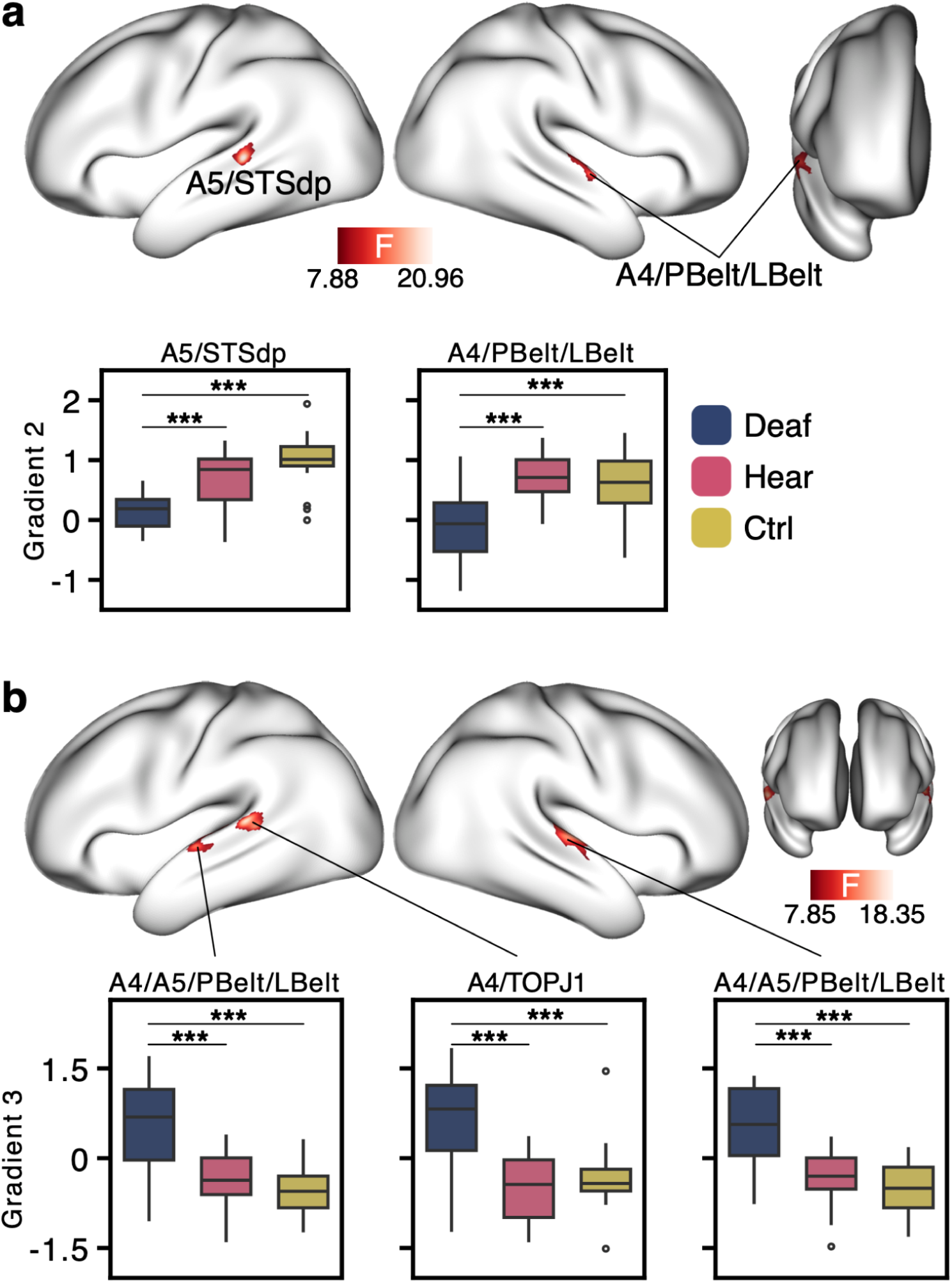
Significant between-group differences of functional gradients. **a)** Significant between-group differences of Gradient 2. b) Significant between-group differences of Gradient 3. Deaf, early deaf with knowledge of cued speech; Hear, hearing controls with knowledge of cues speech; Ctrl, hearing controls without knowledge of cued speech.

**Figure 7.**
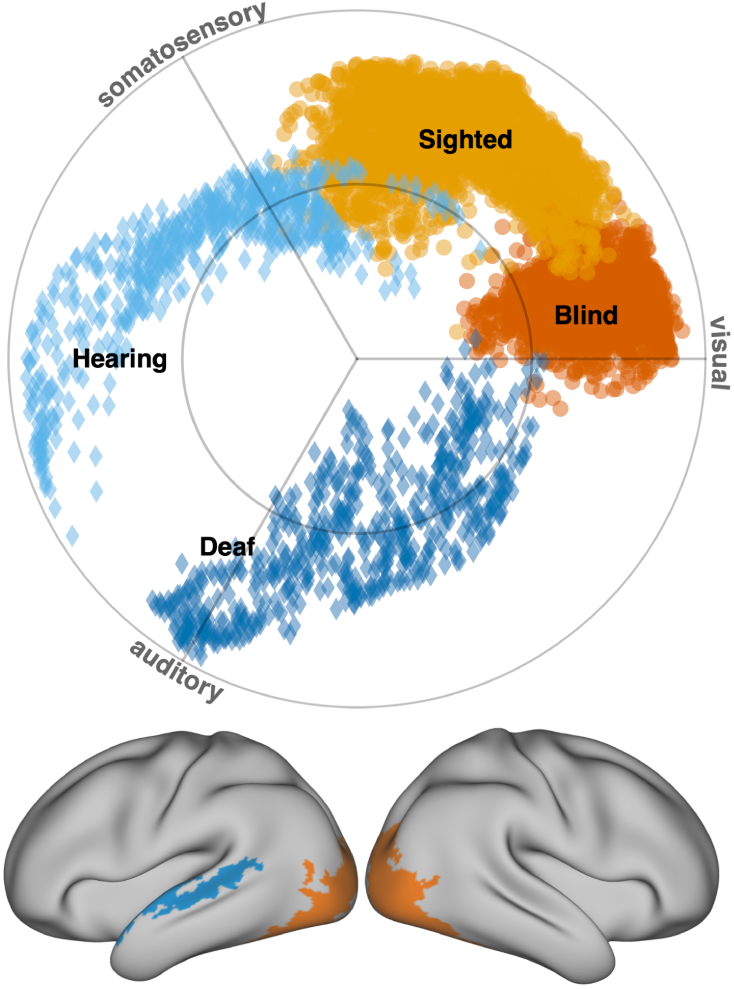
Sensory integration difference between sensory deprived individuals and controls. Early sensory deprived individuals exhibited the most significant sensory integration change in bilateral extrastriate cortex (blindness, orange color) and left superior temporal cortex (deafness, blue color). Their sensory integration patterns are shown in the polar plot.

Consistent with the connectivity results, all significant between-group differences in functional gradients were driven solely by the distinction between early deaf individuals (Deaf) and hearing controls (Hear and Ctrl). Notably, neither traditional functional connectivity analysis nor functional gradient mapping detected differences between the two hearing groups (Hear and Ctrl). In contrast, sensory integration mapping successfully identified robust differences associated with language experience. This dissociation highlights the superior sensitivity of the sensory integration framework in detecting subtle functional plasticity that is not observable using conventional macroscale mapping techniques.

## Discussion

In this study, we demonstrate that early sensory deprivation drives targeted reorganization at the boundaries of deprived primary sensory cortex, while preserving the overarching hierarchical architecture of the brain. By applying a function-based sensory integration framework to cohorts of early blind and early deaf individuals, we identified distinct hotspots of reorganization: the extrastriate cortex in early blindness and the superior temporal cortex in early deafness. These localized alterations were independently validated by functional gradient analysis, confirming their consistency across complementary methodological approaches. In contrast, higher-order association areas maintained stable functional profiles. Furthermore in the early deaf study, the sensory integration framework exhibited unique sensitivity by capturing reorganization of language areas related to expertise in cued-speech, which conventional connectivity and gradient analyses failed to detect. Together, these findings characterize cortical reorganization following early sensory deprivation not as a fundamental remapping of the system, but as a stable hierarchical scaffold that accommodates localized shifts in sensory integration.

### Localized Reorganization of Sensory Integration Adjacent to Deprived Primary Cortex

In early blind individuals, the extrastriate cortex next to deprived primary visual cortex exhibited the most pronounced alteration in sensory integration relative to sighted controls. This region has long been recognized as a major locus of cross-modal plasticity, supporting auditory (Arno et al., 2001; Collignon et al., 2007; Gougoux et al., 2005; Poirier et al., 2006; Voss et al., 2008) and tactile (Amedi et al., 2007; Korczyk et al., 2026; Merabet et al., 2008; Rączy et al., 2019; Sadato et al., 1996; Uhl et al., 1991) processing, as well as higher-order functions including working memory, numerical reasoning, and language (Amedi et al., 2004; Park et al., 2011; Rimmele et al., 2019); (Kanjlia et al., 2016); (Amedi et al., 2007; Bedny et al., 2011; Röder et al., 2002; Watkins et al., 2012). Crucially, our functional gradient analyses revealed prominent group differences in the principal gradients (G1 and G2) within this same territory, spatially converging with the sensory integration results. Together, these findings indicate that cortical plasticity in early blindness is not diffusely distributed across the visual network but is tightly anchored to the extrastriate cortex, adjacent to the primary visual area.

Similarly, in early deaf individuals, sensory-integration differences were most pronounced in the superior temporal cortex. Prior work has consistently identified the superior temporal cortex as a hub for functional recruitment, encompassing visual (Bottari et al., 2014; Emmorey et al., 2011; Fine et al., 2005; Finney et al., 2001, 2003; Karns et al., 2012; Leonard et al., 2012; Zimmermann et al., 2021, 2024) and tactile (Auer et al., 2007; Karns et al., 2012) responses, as well as working-memory engagement (Cardin et al., 2018; Ding et al., 2015). Complementary differences in Gradients 2 and 3 reinforced this localization, confirming that cortical reorganization in early deafness occurs within the belt and parabelt regions adjacent to the primary auditory cortex (A1).

The consistent confinement of reorganization to cortical territories adjacent to deprived primary areas reveals a fundamental organizational constraint on neuroplasticity. These unimodal association areas—such as the extrastriate and superior temporal cortex—occupy an intermediate hierarchical position. They are anatomically poised to integrate sensory information while maintaining privileged access to both primary inputs and higher-order systems. This unique architecture may render them functionally flexible, allowing the cortex to recalibrate sensory integration strategies locally without disrupting the brain’s large-scale topology. In contrast, higher-order association regions, which operate as multimodal convergence hubs, appear to preserve their canonical integrative roles. Evidence from developmental animal models similarly indicates that early sensory deprivation can induce substantial cross-modal reorganization within sensory cortex while respecting broader architectural constraints, including the emergence of novel cortical territories or altered areal boundaries within the visual system (Kahn & Krubitzer, 2002; Rakic et al., 1991). Consistent with this interpretation, primate models of congenital blindness have shown that early deprivation induces local reorganization within mid-level cortical stages while preserving the overarching hierarchical organization (Magrou et al., 2018). This pattern suggests that cortical plasticity preferentially unfolds within these transitional zones, enabling functional compensation without perturbing the brain’s global architecture.

### Feedback dominance shapes intrinsic occipital dynamics after sensory loss

Enhanced intrinsic connectivity between the extrastriate cortex and higher-order association regions emerged as a key finding in early blindness (Fig. 2b). These regions—including fronto-parietal and default mode areas supporting language, numerical reasoning, and working memory—have been repeatedly shown to engage the occipital cortex during cognitive tasks (Amedi et al., 2003, 2004; Bedny et al., 2011; Bonino et al., 2008; Kanjlia et al., 2016; Park et al., 2011; Rimmele et al., 2019; Röder et al., 2002; Watkins et al., 2012, Abboud & Cohen, 2019). Consistent with these task-based findings, resting-state studies have revealed strengthened coupling between visual and language-related fronto-temporal regions, as well as areas involved in social and semantic processing, in blind individuals (Bedny et al., 2011; Burton et al., 2014; Butt et al., 2013; Deen et al., 2015; Liu et al., 2007; Sabbah et al., 2016; Striem-Amit et al., 2015; Watkins et al., 2012), suggesting that the intrinsic activity of the deprived visual cortex becomes increasingly modulated by higher-order networks. This pattern may reflect the network-level substrate that supports higher-order recruitment during active processing (Pelland et al., 2017), rather than direct evidence of such recruitment under resting conditions. In the absence of feedforward sensory information, we propose that this pattern reflects a dominance of feedback processing, where top-down signals from association cortex act as the primary driver of spontaneous activity in the visually deprived occipital lobe.

This mechanism could also help resolve the apparent conflicting results between task-based cross-modal sensory recruitment and resting-state connectivity. While task-based neuroimaging consistently demonstrates auditory and tactile activations in the occipital cortex of early blind individuals (Amedi et al., 2007; Arno et al., 2001; Collignon et al., 2007; Gougoux et al., 2005; Merabet et al., 2008; Poirier et al., 2006; Sadato et al., 1996; Uhl et al., 1991; Voss et al., 2008), resting-state analyses—including our own—often reveal decreased functional connectivity between the auditory and visual areas (Burton et al., 2014). This dissociation is grounded in neuroanatomy: studies in nonhuman primates reveal sparse direct projections from primary auditory cortex (A1) to V1, but extensive feedback pathways from transmodal association areas to the visual cortex (Beer et al., 2011; Rockland & Ojima, 2003). Consequently, in early blindness, the occipital cortex might receive predominantly feedback signals from higher-order regions rather than feedforward sensory input, as occurs during typical development. This anatomical redirection likely shapes intrinsic activity of the visual cortex, thereby facilitating the recruitment of visual areas for higher-order cognitive functions via top-down pathways.

To systematically assess functional reorganization following early sensory deprivation, naturalistic paradigms offer a promising approach. Comparing sensory integration and functional gradients across rest and movie-watching conditions reveals pronounced modulations in both visual and auditory areas (Samara et al., 2023; Wei et al., 2024). Task-based sensory deprivation studies also underscore the necessity of external sensory stimuli to elicit cross-modal activations. Recent studies employing naturalistic stimuli further demonstrated that, in early blind individuals, the extrastriate cortex shows stronger coupling with speech features than in sighted controls, whereas in early deaf individuals, the superior temporal cortex exhibits enhanced coupling with visual features (Orsenigo et al., 2025). Consistent with this pattern, the secondary auditory areas in deaf individuals have been shown to respond to high-level visual information during a naturalistic animated movie, suggesting that these regions may support the processing of rich non-verbal stimuli (Zimmermann et al., 2024). These effects were accompanied by strengthened connectivity between sensory and higher-order networks, reinforcing the view that resting-state dynamics capture only the stable backbone of reorganization, while naturalistic contexts unveil its functional expressivity. Together, these findings highlight that cortical plasticity following early sensory deprivation unfolds along two complementary axes—local cross-modal adaptation and global higher-order recruitment—whose relative prominence depends on behavioral context.

### Sensory integration reveals subtle functional reorganization beyond connectivity measures

The present findings establish the sensory integration framework as a highly sensitive tool for detecting functional reorganization associated with perturbations of the sensory systems. This advantage was most evident in the early deaf cohort. While traditional functional connectivity and gradient analyses successfully identified the robust, deprivation-driven alterations in the superior temporal cortex, they failed to detect the more subtle plasticity associated with language experience. In contrast, the sensory integration model uniquely uncovered functional alterations within the left inferior frontal gyrus and frontopolar cortex—regions where reorganization was driven by the specific modality of language acquisition (Cued Speech) rather than sensory deprivation itself. This pattern is consistent with previous studies of cued-speech users showing enhanced recruitment of superior temporal and inferior frontal language-related regions during multimodal speech processing (Sarré and Cohen 2025), as well as evidence that early language experience shapes the organization and lateralization of higher-order language systems in deaf individuals (Leybaert and D’Hondt 2003). The involvement of the left IFG is particularly relevant given its central role in phonological and semantic language processing (Poldrack et al. 1999), as well as its strong functional coupling with superior temporal cortex (Bokde et al. 2001; Saur et al. 2008; Friederici 2011). Similarly, altered sensory integration within the frontopolar cortex may reflect adaptive changes in high-level integrative and cognitive control systems engaged during visually guided language processing. Together, these findings suggest that experience-dependent plasticity propagates preferentially along existing auditory-language and higher-order associative pathways, rather than reflecting unrestricted cortical reassignment.

The heightened sensitivity of this analytic approach arises from the intrinsic design of the sensory integration framework. Unlike connectivity or gradient analyses, which rely on extrinsic patterns of pairwise correlations or shared variance across the whole brain, our model decomposes the local neural signal into biologically grounded sensory axes. This approach allows for the quantification of intrinsic processing strategies—specifically, how cortical regions rebalance sensory dominance—independent of its long-range coupling strength. By anchoring cortical organization to a common sensory reference space, the framework provides a direct index of how cortical territories recalibrate their functional affiliations. This capability makes it uniquely suited to disentangle the overlapping effects of sensory deprivation and cognitive experience, such as language learning, on brain organization.

## Limitations

Several limitations should be considered. While our results suggest that reorganized sensory areas are primed for cross-modal or higher-order processing, future studies employing naturalistic or specific cognitive tasks are needed to confirm how these intrinsic shifts in sensory integration strategies are deployed during real-time processing. Second, although we successfully dissociated the effects of sensory deprivation from language experience (Cued Speech) in the early deaf cohort, this analysis relied on group-level comparisons. We did not include individual behavioral measures of language proficiency or cross-modal discrimination. Consequently, linking the specific magnitude of the observed angular shifts to individual differences in behavioral adaptation remains a critical avenue for future research. Finally, as a cross-sectional study, our findings represent the endpoint of adult plasticity. The hierarchical structure might be different if the deprivation occurs later in childhood after sensory pathways are already established. Longitudinal studies tracking cortical development in sensory-deprived infants would be required to map the precise developmental trajectory, specifically to determine how the conservation of the global hierarchy interacts with the emergence of local reorganization during critical periods of brain maturation.

## Conclusion

In conclusion, our findings converge on a unifying principle: regions adjacent to deprived primary cortex serve as local hubs of reorganization, preserving the brain’s large-scale architecture while enabling flexible adaptation to sensory loss. In addition to identifying where plasticity occurs, the sensory integration framework captures how it unfolds through shifts in the balance among sensory systems. Its heightened sensitivity to subtle functional changes across blindness and deafness underscores its potential as a generalizable approach for mapping cortical reorganization and for probing the adaptive limits of human neuroplasticity across the lifespan and in both health and disease.

## Acknowledgements

This research has received funding from the European Research Council (ERC) under the European Union’s Horizon 2020 research and innovation programme (Grant agreement No. 866533) awarded to D.S.M. It was supported by the NIHR Oxford Health Biomedical Research Centre (NIHR203316). The views expressed are those of the authors and do not necessarily reflect those of the NIHR or the Department of Health and Social Care. The Wellcome Centre for Integrative Neuroimaging was supported by core funding from the Wellcome Trust (203139/Z/16/Z and 203139/A/16/Z). We also thank the financial support from the program of China Scholarship Council (CSC) awarded to W.W. (File No. 202108330054).

