## Supplementary materials for "Early sensory deprivation drives local reorganization of sensory integration within a conserved global hierarchy"

### Preprocessing

##### Preprocessing of the early blind datasets

Each T1-weighted (T1w) image was reoriented to left–posterior–inferior (LPI) orientation and corrected for intensity non-uniformity using N4BiasFieldCorrection (Tustison et al., 2010). Skull stripping was performed on the bias-corrected images, followed by tissue segmentation using FSL FAST (Zhang et al., 2001). Cortical surface reconstruction was conducted using FastSurfer (Henschel et al., 2020).

For the functional data, the first 10 volumes of each resting-state fMRI run were discarded to allow for signal stabilization. Images were reoriented to LPI orientation and motion-corrected by aligning all volumes to the mean functional image. A high-pass temporal filter (cutoff = 0.01 Hz) was applied to remove low-frequency drifts.

Functional volumes were averaged and registered to the native anatomical space using boundary-based registration (Greve & Fischl, 2009). The time series were then projected onto individual cortical surfaces using trilinear interpolation and resampled to the HCP fsLR_32k surface mesh (59,412 vertices excluding the medial wall).

Nuisance regression included signals from white matter and cerebrospinal fluid, as well as motion-related spike regressors. To account for superficial cortical regions not covered during fMRI acquisition, a mutual brain mask was applied to exclude unsampled areas from subsequent analyses (Supplementary Fig. 1a). Finally, surface-based spatial smoothing was applied within the mutual mask using a 4 mm FWHM kernel via the cifti-smoothing command in Connectome Workbench (Glasser et al., 2013).

##### Preprocessing of the early deaf dataset

The T1w images exhibited pronounced intensity non-uniformity artifacts that could not be fully corrected by the default fMRIPrep workflow. Therefore, an initial bias field correction was applied using FSL FAST prior to preprocessing. The images then underwent a second intensity non-uniformity correction using N4BiasFieldCorrection, and the resulting bias-corrected images were used as the anatomical reference throughout fMRIPrep processing.

Skull stripping and tissue segmentation were performed using FSL FAST, and cortical surface reconstruction was carried out using FreeSurfer recon-all (Dale et al., 1999). Nonlinear spatial normalization to the MNI152NLin6Asym template was performed using ANTs.

Each subject completed three multi-echo resting-state fMRI runs. For each run, the first five dummy volumes were removed. A reference volume was generated from the shortest echo and used for motion correction. Motion parameters were estimated relative to this reference, and all volumes were realigned accordingly. Functional images were coregistered to the T1w reference using boundary-based registration.

The preprocessed BOLD time series were projected onto the cortical surface and resampled to the HCP fsLR_32k mesh using Connectome Workbench. Due to sampling failures primarily around the medial wall, a mutual surface mask was applied to exclude unsampled regions from further analysis (Supplementary Fig. 1b).

Nuisance regression included 24 Friston motion parameters and the first five aCompCor components extracted from a combined white matter and cerebrospinal fluid mask. The cleaned time series were bandpass filtered (0.01–0.08 Hz) and spatially smoothed within the mutual mask using a 6 mm FWHM kernel.

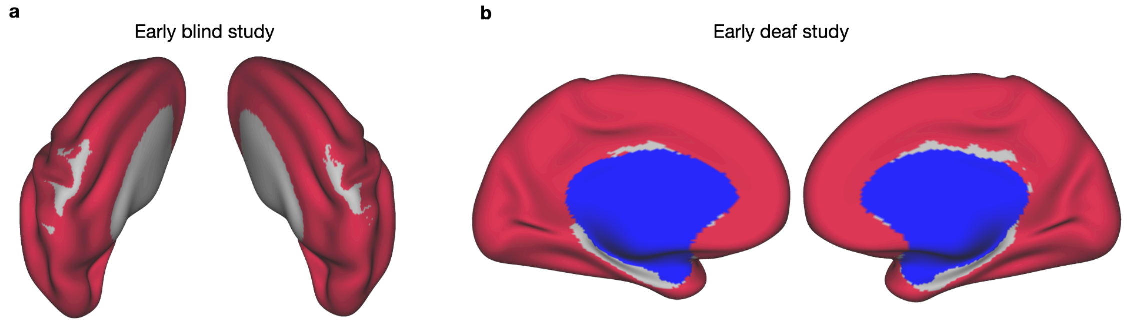


**Supplementary Figure 1. Mutual mask for early blind and early deaf study.** a) Mutual mask for early blind datasets. Areas covered by the mutual mask were labeled by the red color. b) Mutual mask for early dead dataset. Areas covered by the mask were labeled by the red color. Bilateral medial walls were labeled by the blue color.

#### Colinearity

To map sensory integration, we quantified the contributions of different sensory modalities to the fMRI signal at each vertex using a general linear model (GLM). The averaged time series from the primary visual cortex (V1), primary somatosensory cortex (S1), and primary auditory cortex (A1) served as predictors for the time series at each vertex. Collinearity among these primary sensory signals can inflate the variance and standard error of the regression coefficients, potentially reducing the reliability of our sensory integration model. To assess collinearity, we calculated the Variance Inflation Factor (VIF) for each predictor, specifically the averaged time series from V1, S1, and A1, for each subject.

VIF quantifies the extent to which the variance of a regression coefficient is increased due to collinearity. For each predictor in a regression, the VIF is computed by regressing this predictor on all other predictors in a separate regression model. The VIF for a predictor is given by:

$$VIF_i = \frac{1}{1-R_i^2}$$

where $R_i^2$ is the coefficient of determination from the regression of the ith predictor on the other predictors.

Typically, a VIF less than 5 indicates acceptable collinearity in a linear regression. Our results showed that all subjects had low VIFs for the primary sensory signals, indicating minimal collinearity. Therefore, no further orthogonalization procedures were necessary.

**Supplementary Table 1. Variance inflation factors of sensory predictors**

| | Early blind study (n = 98) | | Early deaf study (n = 60) | |
|---|---|---|---|---|
| | mean | std | mean | std |
| TS-V1 | 1.35 | 0.4 | 1.38 | 0.44 |
| TS-S1 | 1.59 | 0.52 | 1.47 | 0.52 |
| TS-A1 | 1.39 | 0.34 | 1.67 | 0.64 |

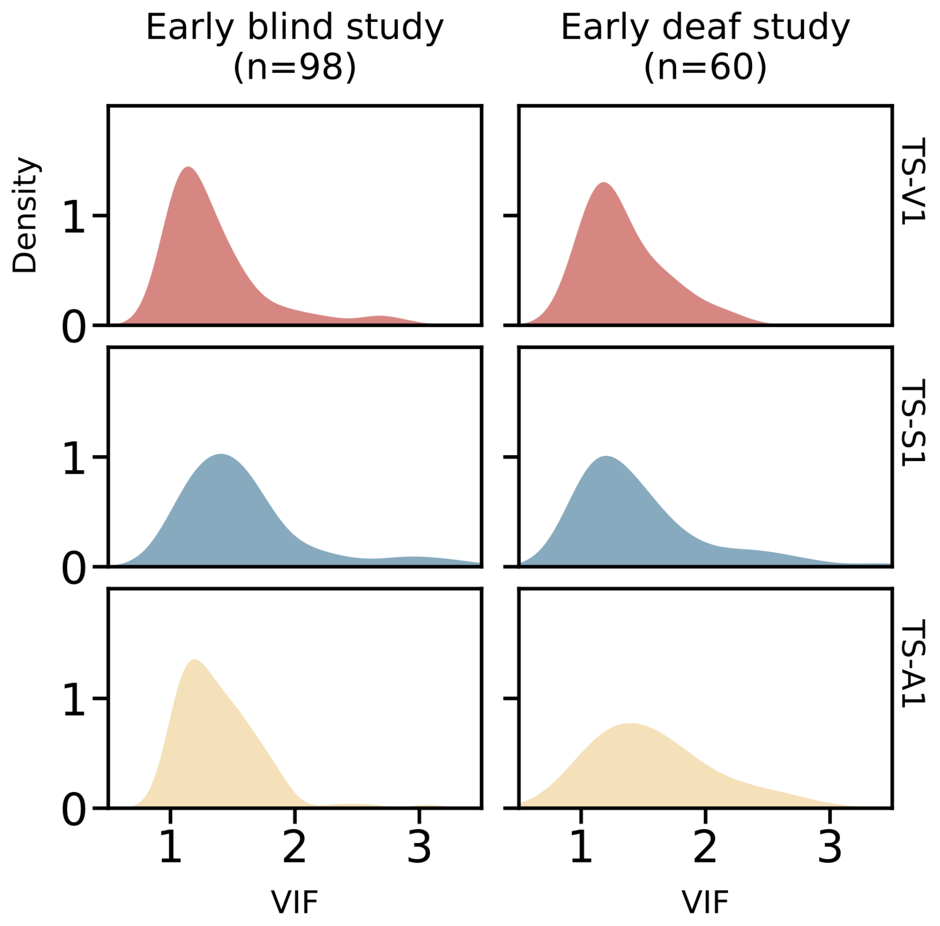


**Supplementary Figure 2. The VIF Distributions of sensory predictors in early blind and early deaf study.** VIF, variance inflation factor; TS, the averaged time series; V1, primary visual cortex; S1, primary somatosensory cortex; A1, primary auditory cortex.

**Supplementary Table 2.** Sensory parameter comparison (early blind vs. sighted control)

| | | visual | | | sensorimotor | | | auditory | | |
|---|---|---|---|---|---|---|---|---|---|---|
| | | **M** | **T** | **P** | **M** | **T** | **P** | **M** | **T** | **P** |
| extra-striate | t | 3.5 | 5.3 | 4.25 | -3.32 | -3.25 | -3.53 | 1.69 | -0.13 | 0.6 |
| | p | 0.001* | 1e-05* | 3.3e-04* | 0.002* | 0.002* | 0.002* | 0.1 | 0.9 | 0.55 |
| V1/V2 | t | 0.83 | 0.91 | 2.98 | -4.43 | -1.26 | -3.26 | 2.44 | 2.29 | 1.8 |

| | | | | | | | | | | |
|---|---|---|---|---|---|---|---|---|---|---|
| | p | 0.42 | 0.37 | 0.007 | 1e-04* | 0.22 | 0.004* | 0.02 | 0.03 | 0.09 |
| V6 | t | 0.03 | 1.37 | 1.43 | -3.06 | -2.08 | -3.81 | 4.71 | 5.37 | 3.01 |
| | p | 0.98 | 0.18 | 0.17 | 0.004* | 0.04 | 0.001* | 4e-05* | 0* | 0.006 |
| STSdp | t | -1.08 | -2.89 | -2.19 | 2.21 | 1.81 | 1.32 | 0.51 | 0.05 | -0.68 |
| | p | 0.29 | 0.006 | 0.04 | 0.03 | 0.08 | 0.2 | 0.61 | 0.96 | 0.5 |

*M, Montreal dataset; T, Trento dataset; P, Paris dataset; * p < 0.0056 (Bonferroni correction for the each cluster-based comparison, 0.05/9)*

#### GCCA-based gradient

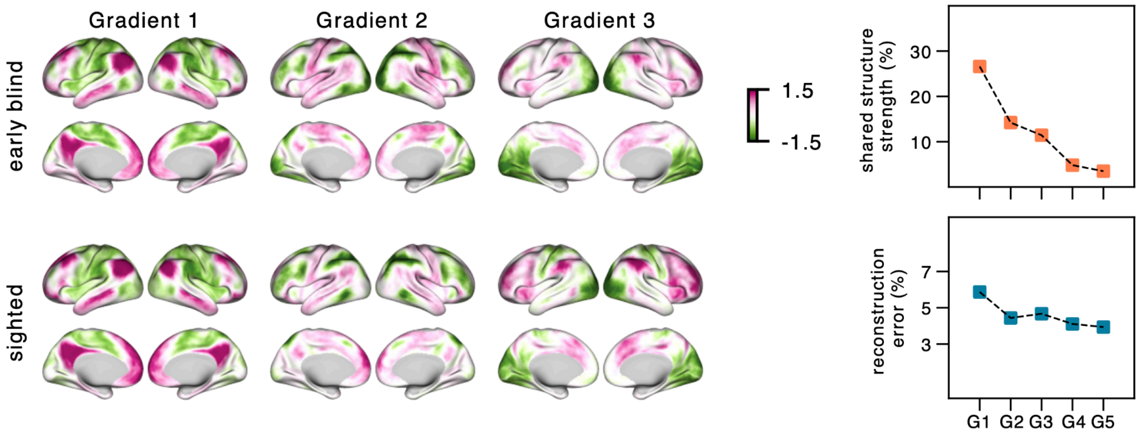


**Supplementary Figure 3. GCCA-based gradients of early blind and sighted control.** The first three GCCA-based gradients were averaged across all datasets and illustrated on the cortical surface. As GCCA captures the common features across all the subjects, the shared structure strength of the first five gradients was shown in the top right plot. The reconstruction error or variance explained by each gradient was displayed in the bottom right plot.

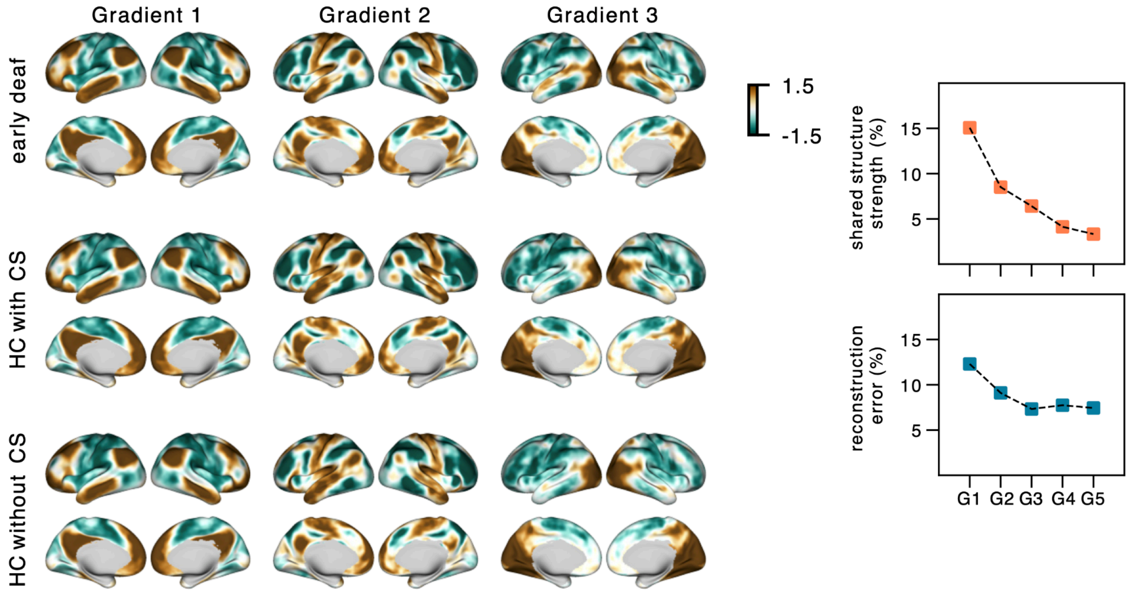


**Supplementary Figure 4. GCCA-based gradients of the early deaf dataset.** The group-averaged GCCA-based gradients were illustrated on the cortical surface. As GCCA captures the common features across all the subjects, the shared structure strength of the first five gradients was shown in the top right plot. The reconstruction error or variance explained by each gradient was displayed in the bottom right plot.
